# Interaction of BIR2/3 of XIAP with E2F1/Sp1 Activates MMP2 and Bladder Cancer Invasion by Inhibiting Src Translation

**DOI:** 10.1101/309765

**Authors:** Jiheng Xu, Honglei Jin, Jingxia Li, Junlan Zhu, Xiaohui Hua, Zhongxian Tian, Maowen Huang, Rui Yang, Haishan Huang, Chuanshu Huang

**Affiliations:** School of Laboratory Medicine and Life Science, Wenzhou Medical University, Wenzhou, Zhejiang, 325035, China; Nelson Institute of Environmental Medicine, New York University School of Medicine, Tuxedo, NY 10987, USA

**Author notes:** Corresponding author: Haishan Huang, Ph.D. School of Laboratory Medicine and Life Science, Wenzhou Medical University, Wenzhou, Zhejiang, China 325035.

**Keywords:** BIR domains of XIAP, Invasion, E2F1, Sp1, Src

## Abstract

Although X-linked inhibitor of apoptosis protein (XIAP) is associated with cancer cell behaviors, the structure-based function of XIAP in promotion human bladder cancer (BC) invasion is barely explored. Herein, we discovered that ectopic expression of the BIR domains of XIAP rescued the MMP2 activation and invasion in XIAP-deleted BC cells, while Src was further defined as a XIAP downstream negative regulator for MMP2 activation and BC invasion. The inhibition of Src expression by BIR domains was caused by attenuation of Src protein translation upon miR-203 upregulation resulting from direct interaction of BIR2 and BIR3 with E2F1 and Sp1, consequently leading to fully activation of E2F1/Sp1. Our findings provide a novel insight into understanding of specific function of BIR2 and BIR3 of XIAP in BC invasion, which will be highly significant for the design/synthesis of new BIR2/BIR3-based compounds for invasive BC treatment.

## Introduction

X-linked inhibitor of apoptosis (XIAP) is an IAP protein family member and a well-defined inhibitor of the caspase/apoptosis pathway [1-3]. Emerging evidence has revealed that abnormal expression of XIAP is associated with tumorigenesis in breast cancer [4], prostate cancer [5-7], acute and chronic leukemia [8-10], bladder cancer [11] and other types of cancers [12,13]. It is notable that XIAP overexpression is particularly associated with the progression and aggression of malignant cancer [14,15]. Thus, XIAP is widely considered as an important player in cancer development and malignancy behaviors.

There are four functional domains in XIAP: three repeats of the baculovirus IAP repeat (BIR) domain at its NH_2_ terminus and a RING finger domain near its COOH terminus. The formers are mainly responsible for its anti-apoptotic function by inhibiting caspase-3, −7 and −9, and the latter contains E3 ubiquitin ligase activity, allows IAPs to ubiquitinize themselves, caspase-3, and caspase-7 *via* the proteasome-dependent mechanism [16]. The studies from our laboratory and others reveal novel functions of XIAP beyond anti-apoptotic function [17-19]. For example, XIAP upregulates cyclin D1 expression *via* an E3 ligase-mediated protein phosphatase 2A/c-Jun axis [20] and upregulates cyclin E expression as a result of direct binding of E2F1 by the BIR domains, which promotes human colon cancer cell growth [21]. XIAP also enhances human invasive BC cell proliferation due to the BIR domain-mediated *c-Jun/miR-200a/EGFR* axis [22]. The RING domain of XIAP interacts with RhoGDP dissociation inhibitor α protein to inhibit RhoGDIα SUMOylation at Lys-138, subsequently affecting human colon cancer cell motility [23,24]. Moreover, downregulation of the tumor suppressor p63α protein expression by the RING domain of XIAP promotes malignant transformation of bladder epithelial cells [25]. Thus, although XIAP was originally classified as an inhibitor of apoptosis protein family member, the function of XIAP on cancer cell proliferation, motility and transformation and its signaling pathway have attracted a great deal of attention.

Our recent preliminary study emphasizes the novel role of XIAP on BC cancer invasion and reveals that XIAP promotes bladder cancer invasion through its BIR domains, indicating a previously underappreciated role of BIR domains to promote invasive activity of cancer cells. Thus, we further dissected the signaling pathways related to this important function in the current study. We discovered that this novel function is mediated by specific activating MMP2 due to BIR domain-initiated suppression of Src protein translation. Moreover, the BIR domains of XIAP attenuated Src protein translation due to directly interaction of BIR2 and BIR3 with E2F1 and Sp1, respectively, leading to miR-203 transcription, and its binding to Src mRNA 3’-UTR region.

## Results

### XIAP BIR domains specifically promoted MMP2 activation and BC invasion in human BC cells

XIAP contains three repeat BIR domains in the N terminus and one RING domain in the C terminus as schematically shown in Figure 1A. To uncover the potential role of these domains in mediation of human high invasive BC behaviors, we employed two human invasive BC cancer cell lines, UMUC3 and T24T. As shown in Figure 1B, inhibition of XIAP expression by its specific shRNA resulted in a profound specific decrease in MMP2 activation without affecting pro-MMP2 protein in T24T cells. Unexpectedly, the reduction of MMP2 activation was further reversed by ectopic expression of HA-ΔRING (XIAP in the absence of all three BIR domains), but not reversed by ectopic expression of HA-ΔBIR (XIAP in the absence of RING domain). A similar result was also observed in UMUC3 cells (Figure 1C). These findings indicate that the BIR domains but not the RING domain were crucial for XIAP-mediated MMP2 activation in human BC cells.

**Figure 1.**
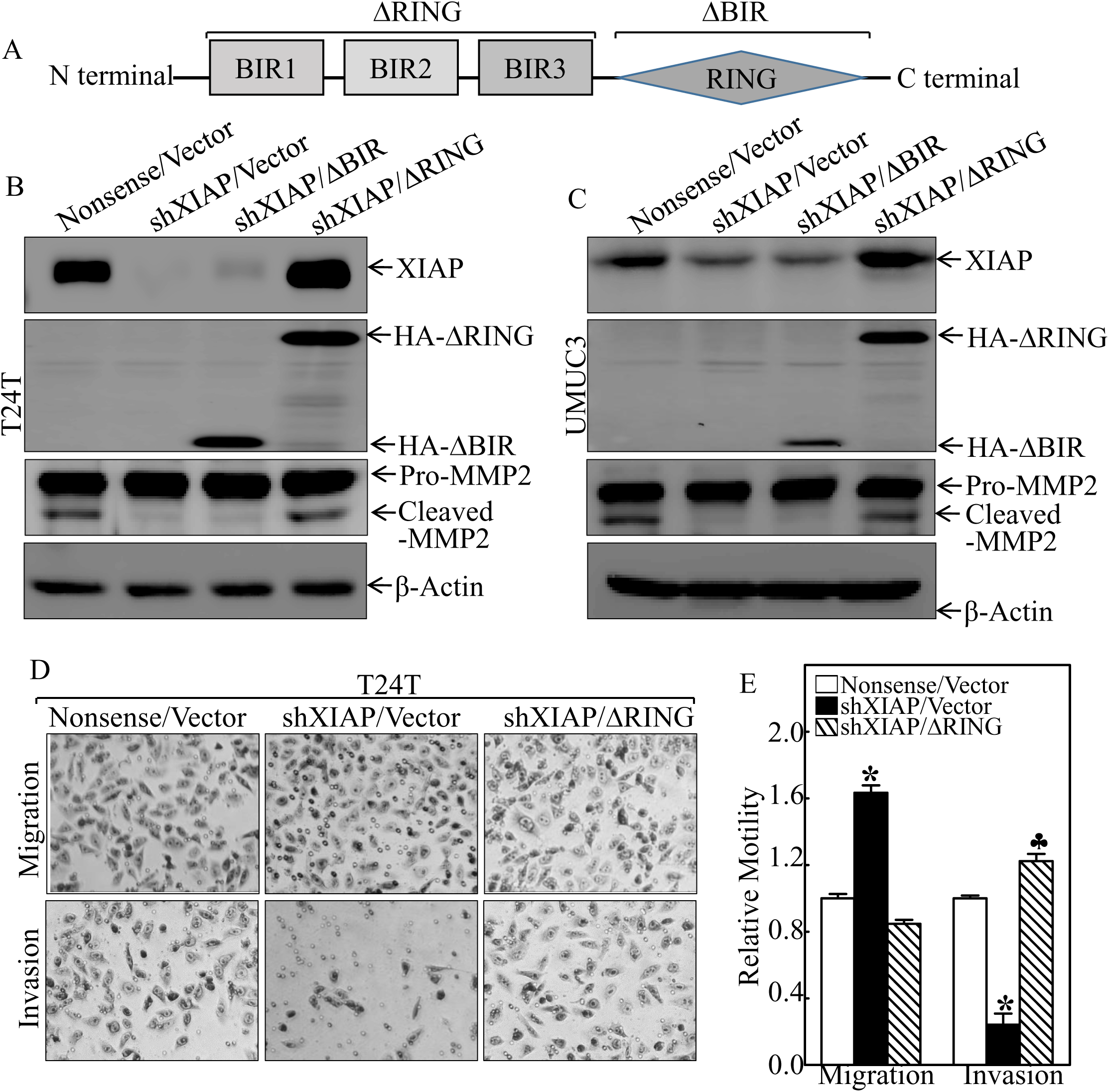
XIAP BIR domains promoted MMP2 activation and BC invasion. **(A)** The schematic structure of XIAP domains. **(B and C)** The indicated cell extracts were subjected to Western Blot for determination of the expression of XIAP and pro-MMP2 and cleaved-MMP2 (activated-MMP2). β-Actin was used as the protein loading control. **(D and E)** T24T (Nonsense/Vector), T24T (shXIAP/Vector), T24T (shXIAPMRING) cells were cultured in uncoated chambers or pre-coated Matrigel chambers for 24 hours. The cells were then fixed and stained. The invasion and migration rates were quantified by counting the relative migratory (Transwell) and invasive cells in at least five random fields under a light microscope, and then, the cell numbers were normalized with the insert control according to the manufacturer’s instructions. The bars show the mean±SD of 3 independent experiments. The symbol (*) indicates a significant difference as compared with the vehicle control (p < 0.05), and the symbol (♣) indicates a significant difference compared with T24T (shXIAP/Vector) cells (p < 0.05).

MMP2 degrades cellular matrix components and the basement membrane, and therefore reduces the barriers for cancer cell migration and/or invasion [26]. Therefore, we next determine the capacity of cell migration and invasion of XIAP BIR domains in T24T cells. As shown in Figures 1D and 1E, inhibition of XIAP expression dramatically reduced BC cell invasion, which is consistent with the reduced activated MMP2 level (Figures 1B and 1C). The reduction of BC cell invasion was restored when ectopic expression of ΔRING (Figures 1D and 1E), indicating that BIR domains are crucial for XIAP-mediated BC invasion. Interestingly, inhibition of XIAP expression increased cell migration (Figures 1D and 1E), suggesting that although cancer cell invasion and migration are appealingly linked in many experimental system, but may be divergent in the significance and mechanism in human BC cells as shown in our recently studies [27].

### Src tyrosine kinase protein expression was inhibited by XIAP BIR domains in BC cells and was downregulated in human and mouse BCs

It has been reported that decreased Src protein expression is associated with late-stage bladder tumor progression [28]. Thus, we tested whether different domains of XIAP could regulate the expression of Src in BC cells. As shown in Figure 2A, attenuation of XIAP expression resulted in a profound increase in Src protein expression and this augmentation on Src protein expression was reversed by ectopic expression of HA-ΔRING, but not by HA-ΔBIR. This result indicates that the BIR domains but not the RING domain are crucial for XIAP inhibition of Src protein expression.

**Figure 2.**
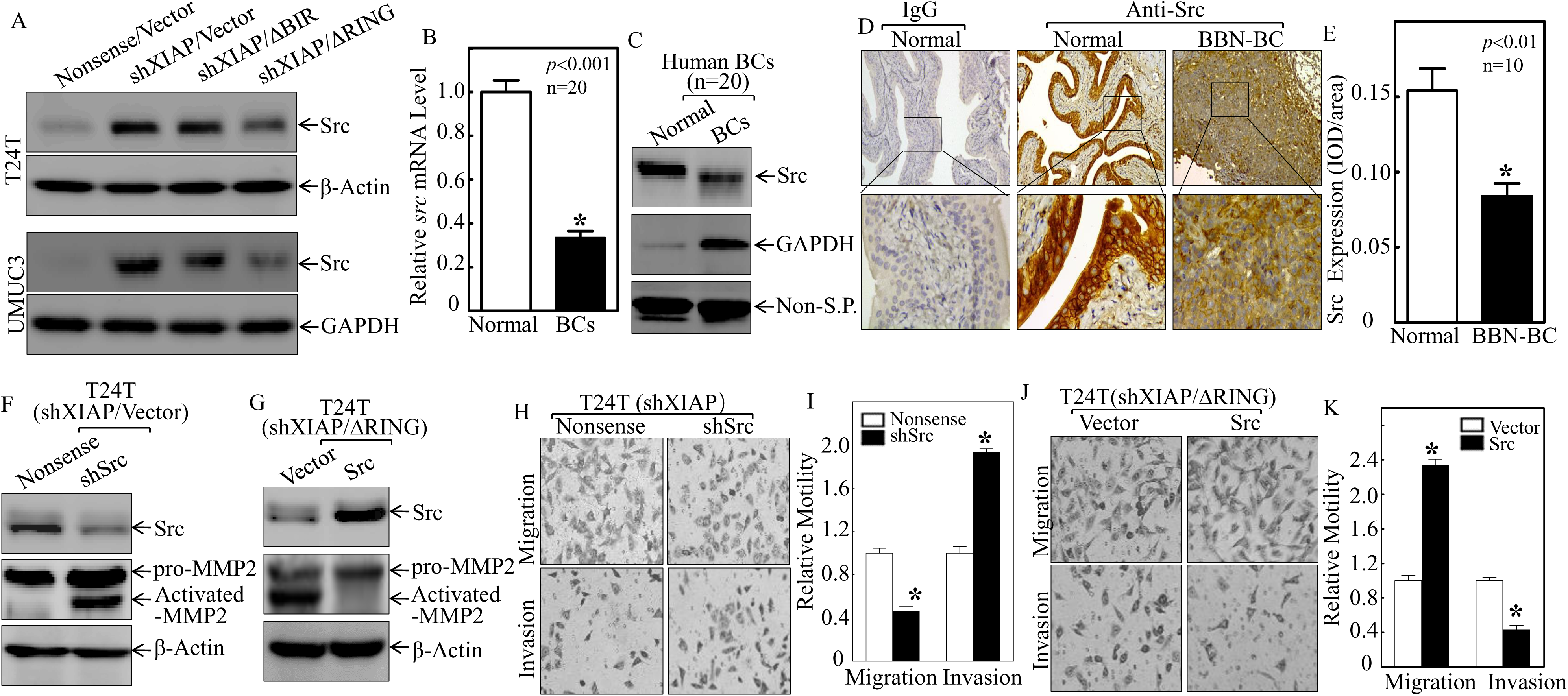
Src downregulation by XIAP BIR domains resulted in BC invasion in human BCs. **(A)** The indicated cell extracts were subjected to Western blot for determination of Src expression. **(B and C)** Total RNA and protein lysates were prepared from human normal and the paired human bladder urothelial cell mixtures separately from 20 patients diagnosed with bladder cancer and subjected to qRT-PCR and Western blotting analyses to determine Src mRNA (B) and protein (C) expression profiles, respectively. (Non-S.P: Non-Specific Protein) **(D and E)** IHC-P was carried out to evaluate Src protein expression in mouse BC induced through consistent exposure of mice to BBN for 20 weeks. The optical density was analyzed as described in “materials and methods”. The symbol (*) indicates a significant decrease in comparison to normal mice (p < 0.01). **(F and G)** The cell extracts from the indicated stable transfectants were subjected to Western Bot for determination of related protein expression. **(H-K)** The indicated stably transfectants were subjected to cell migration and invasion assays as described in “materials and methods”.

Next, we corroborated the negative association of Src protein expression and the stage of bladder cancer using both clinical samples and an *in vivo* animal model. Thus, we evaluated Src expression in 20 pairs of human BC tissues and their adjacent appealingly normal bladder tissues that had been surgically removed from patients diagnosed with BCs. As shown in Figure 2B, a profound reduction in src mRNA expression was observed in human BC tissues, with an overall average of a 3-fold lower relative src mRNA level in comparison with the normal controls. Consistent with the mRNA expression results, significantly decreased Src protein expression was also observed in human invasive bladder cancer tissues (Figure 2C). Moreover, we determined the expression of Src in a mouse model of bladder cancer via consistent exposure of mice to N-butyl-N-(4-hydroxybutyl)-nitrosamine (BBN) for 20 weeks. As illustrated in Figures 2D and 2E, the results from the immunohistochemistry (IHC) staining revealed that Src expression was markedly decreased in mouse invasive BC tissues in comparison to normal mouse bladder tissues. Thus, the reduced expression of Src in invasive bladder cancer and its increase following XIAP depletion indicate the negative correlation between Src and XIAP expression in BCs.

### XIAP BIR domains-mediated Src downregulation was critical for BC cell invasion

To test association of Src suppression by BIR domain with BC invasion, Src was either knocked down or overexpressed in T24T deficient cells, T24T (shXIAP). As shown in Figures 2F and 2G, knockdown of Src in T24T (shXIAP) cells resulted in a greater invasion ability as compared with its nonsense transfectant (Figures 2H & 2I), indicating that Src is a XIAP downstream target and its downregulation is responsible for the XIAP-mediated BC cell invasion. Expectedly, overexpression of Src in T24T (ΔRING) cells attenuated cell invasion in comparison to scramble vector transfectant (Figures 2J and 2K). Our results reveal that Src suppression participates into the XIAP BIR domains-mediated BC cell invasion.

### XIAP BIR domains inhibited Src protein translation through upregulating miR-203 in human BC cells

Our above results showed that the deletion of XIAP BIR domains increased Src protein expression (Figure 3D), indicating that XIAP BIR domains mediate Src inhibition. Therefore, we next determined whether the XIAP BIR domains have a suppressive effect on Src mRNA expression. The results showed that src mRNA levels were nearly comparable in T24T cells with XIAP knockdown, or XIAP knockdown with either BIR domain overexpression or RING domain overexpression (Figure 3A). Similar results were also observed in UMUC3 cells with similar approaches (Figure 3B). Interestingly, depletion of XIAP expression in T24T cells resulted in a faster degradation of Src protein (Figure 3C), indicating that XIAP plays a role in Src protein stabilization, further suggesting that XIAP might regulate Src protein translation. The results from incorporation of 35S-methionine/ cysteine into newly synthesized Src protein in T24T cells in XIAP knockdown cells was markedly increased in comparison to control T24T(nonsense) cells (Figure 3D), further revealing that XIAP did inhibit Src protein translation. Finally, we found that ribosomal S6 appears not related to XIAP-mediated suppression of Src protein translation.

**Figure 3.**
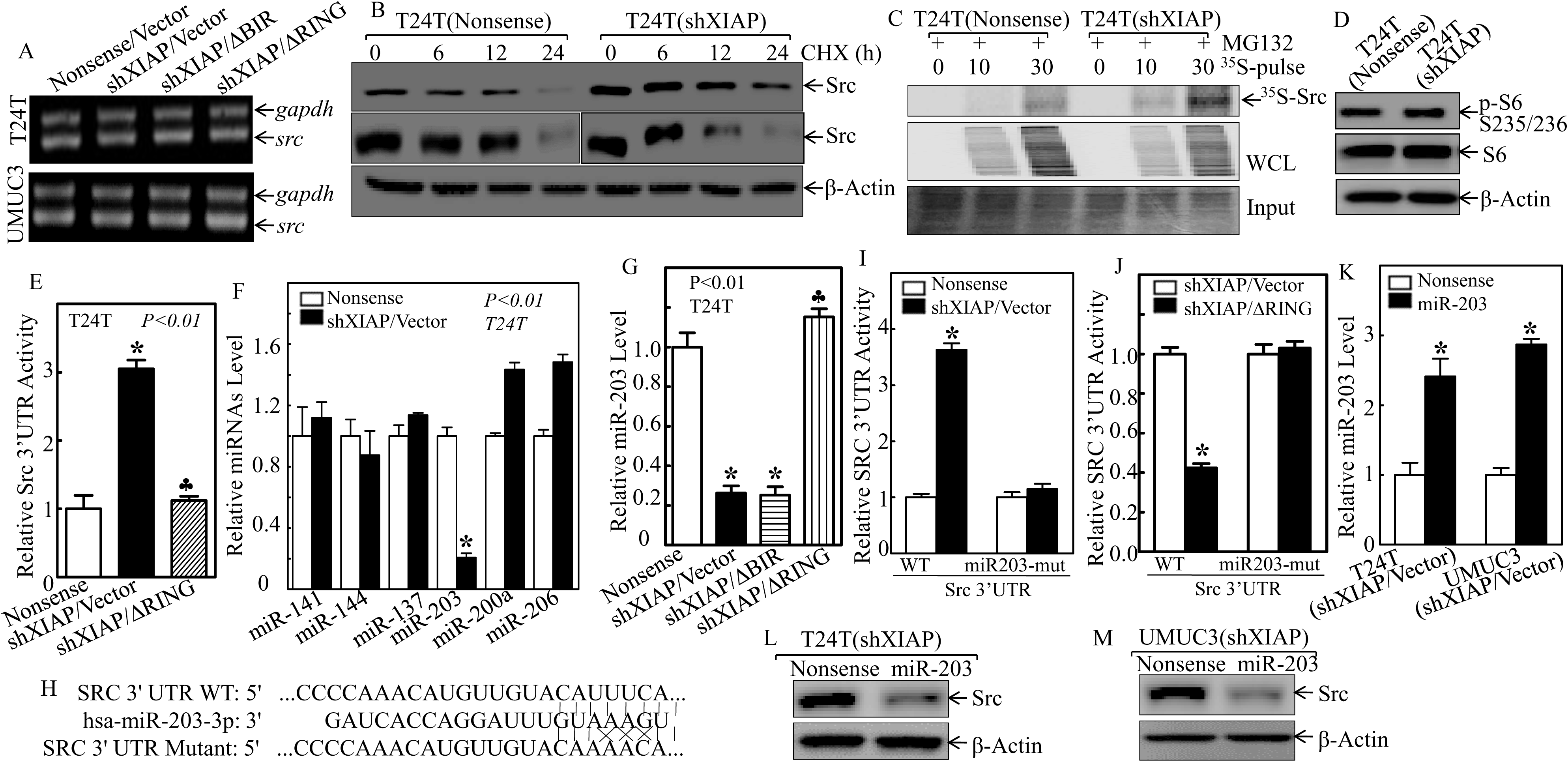
BIR domains of XIAP promoted miR-203 transcription and in turn inhibited Src tyrosine kinase protein translation in human bladder cancer cells. **(A)** Total RNA was isolated from the indicated cells and then subjected to RT-PCR analysis of src mRNA expression. GAPDH was used as a loading control. **(B)** After pre-treatment with MG132 (10 μΜ) for 6 h, T24T (Nonsense) and T24T (shXIAP) cells were subjected to determination of Src protein degradation in the presence of cycloheximide (CHX) (100 μg/ml). β-Actin was used as a protein loading control. **(C)** After pretreatment with MG132 (10 μΜ) for 30 mins, newly synthesized Src protein in T24T (Nonsense) and T24T (shXIAP) cells was monitored with a pulse assay using ^35^S-labeled methionine/cysteine. WCL stands for whole cell lysate. Coomassie blue staining was used for protein loading control. **(D)** The cell extracts were subjected to Western Blot as indicated. β-Actin was used as a loading control. **(E)** The indicated cells were transiently transfected with a Src 3'UTR luciferase reporter and the luciferase activity of each transfectant was evaluated. The results are presented as Src 3′-UTR activity relative to medium control. The symbol (*) indicates a significant increase as compared with T24T (Nonsense/Vector) (p < 0.01). The symbol (^♣^) indicates a significant decrease as compared with T24T (shXIAP/Vector) cells. **(F)** The levels of the indicated microRNAs were evaluated with quantitative real-time PCR. The symbol (*) indicates a significant decrease compared with control cells as indicated (p < 0.01). **(G)** The level of miR-203 in the indicated cells was evaluated with quantitative real-time PCR. The symbol (*) indicates a significant decrease as compared with nonsense cells as indicated (p < 0.01), while the symbol (^♣^) indicates a significant increase as compared with T24T (shXIAP/Vector) cells. **(H)** Schematic of the construction of the src mRNA 3′-UTR luciferase reporter and its mutants aligned with miR-203. **(I and J)** T24T (Nonsense), T24T (shXIAP) cells and T24T (shXIAP/Vector), T24T (shXIAPMRING) cells were co-transfected with wild-type and mutant src 3′-UTR luciferase reporters and pRL-TK, respectively. The luciferase activity of each transfectant was evaluated, and the results are presented as relative to src 3′-UTR activity. The symbol (*) indicates a significant difference in src 3′-UTR activity (p < 0.01). **(K)** T24T (shXIAP/Vector) and UMUC3 (shXIAP/Vector) cells were stably transfected with constructs of miR-203 or its control vector. miR-203 expression was determined with real-time PCR, and the symbol (*) indicates a significant increase as compared with control nonsense transfectant (p < 0.05). **(L and M)** The indicated cell extracts were subjected to Western blotting, and β-Actin was used as the protein loading control.

To explore the mechanisms underlying XIAP suppression of Src protein translation, the potential effect of XIAP on phosphorylation of S6 ribosomal protein was evaluated, and the results showed that phosphorylation of S6 ribosomal protein was comparable in T24T(Nonsense) and T24T(shXIAP) cells (Figure 3E), excluding the possible involvement of S6 ribosomal protein in XIAP inhibition of Src protein translation. We next texted whether XIAP modulated Src mRNA 3’-UTR activity and the results indicated that XIAP knockdown resulted in the augmentation on Src mRNA 3’-UTR activity, whereas ectopic expression of three BIR domains (HA-ΔRING) reversed an increase in Src mRNA 3’-UTR activity (Figure 3F), suggesting that the BIR domains are required for XIAP inhibition of src mRNA 3’-UTR activity. Since microRNAs (miRNAs) could inhibit protein translation *via* interacting mRNA 3’-UTRs [29], a bioinformatics analysis was conducted and showed that miR-141, miR-144, miR-137, miR-203, miR-200a, and miR-503 are putative miRNAs that can bind to the 3’-UTR region of Src mRNA (Table 1). The results for evaluation of these putative miRNAs indicated that knockdown of XIAP only attenuated miR-203 expression in T24T cells (Figure 3G), and the reduction on miR-203 expression was further reversed by ectopic expression of XIAP in the absence of the BIR domains (Figure 3H), indicating that the BIR domains specifically inhibit miR-203 expression in human BC cells. Moreover, the point mutations of the miR-203 binding site in the Src mRNA 3’-UTR reporter completely abolished the increased luciferase activity due to XIAP knockdown in T24T cells (Figures 3I & 3J), revealing that miR-203 binding site is crucial for XIAP/BIR inhibition of Src mRNA 3’-UTR activity. This notion was great supported by the results obtained in T24T (shXIAP/ΔRING) in comparison to that in T24T (shXIAP/Vector) cells (Figure 3K). These results reveal that miR-203 directly binds to 3’-UTR of Src mRNA and mediates the BIR domains inhibition of Src protein translation. To unravel the role of miR-203 in regulation of Src protein expression, miR-203 was transfected into T24T (shXIAP) and UNUC3 (shXIAP) cells. As shown in figure 3L and 3M, overexpression of miR-203 abolished Src protein expression in both T24T (shXIAP) and UNUC3 (shXIAP) cells, indicating that miR-203 inhibits Src protein expression. Collectively, our study demonstrates that the XIAP BIR domains promote miR-203 expression, resulting in an increase in miR-203 interacting with the src mRNA 3’-UTR and in turn inhibiting Src protein translation.

**Table 1:**
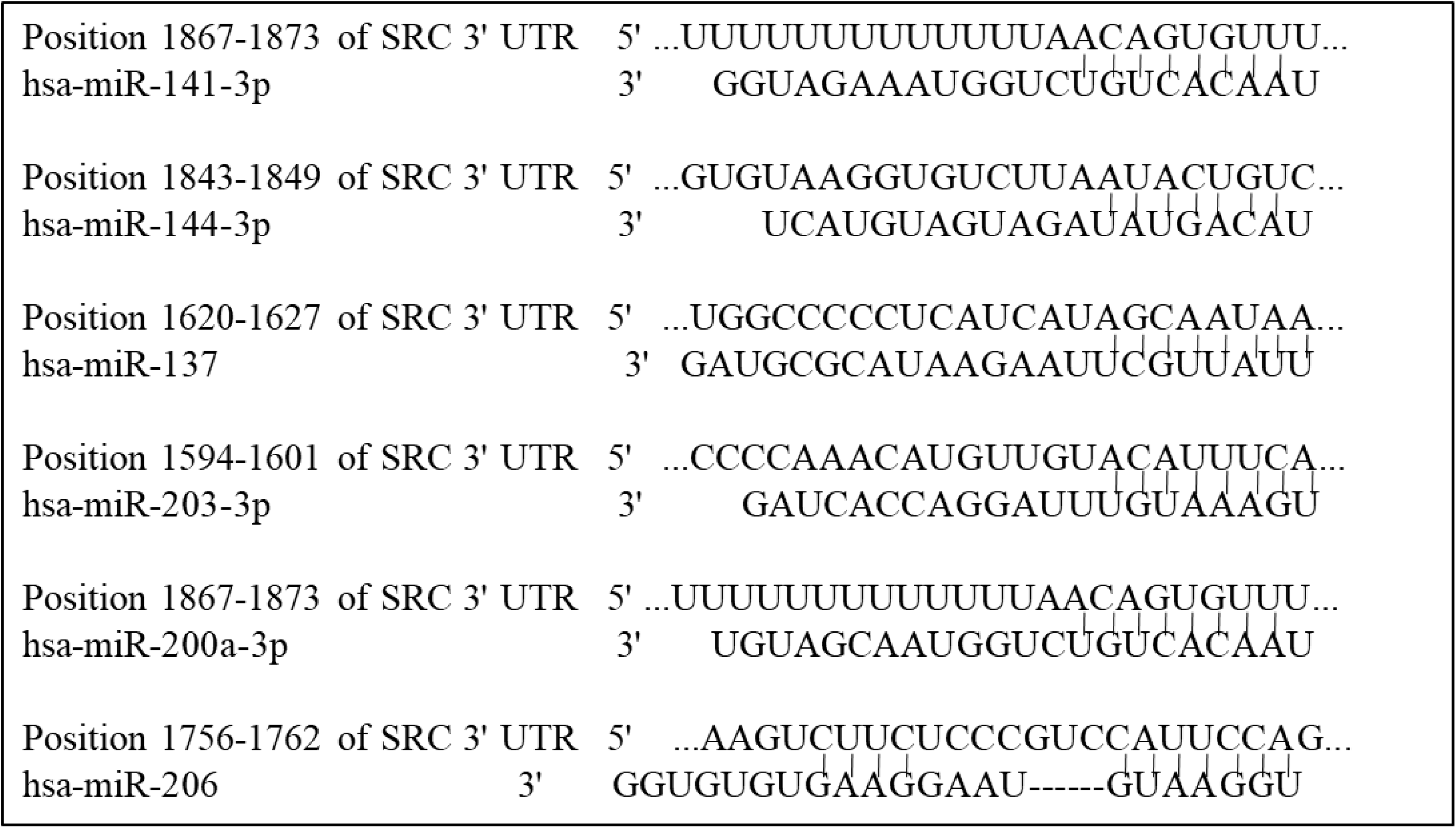
The potential miRNAs binding sites in the Src mRNA 3’UTR region

### XIAP BIR domains promoted miR-203 transcription through activation of E2F1 and Sp1

Since miRNAs possess differential stability in human cells [30], the effect of XIAP/BIR on miR-203 stability were evaluated. As shown in Figure 4A, neither inhibition of XIAP expression nor ectopic expression of HA-ΔRING in T24T (shXIAP) cells showed a significant regulatory effect on miR-203 stability as compared with control transfectants. Pre-miRNAs are regulated at transcription and are processed to mature miRNAs by enzymes, such as dicer and argonaute 2 [31,32]. To examine whether the BIR domains of XIAP regulate miR-203 at transcriptional level, we determined the effect of XIAP and its BIR domains on pre-miR-203 expression as well as its promoter activity. The results showed that both pre-miR-203 abundance and its promoter-driven luciferase reporter activity were impaired in XIAP knockdown cells, whereas ectopic expression of ΔRING domains restored both pre-miR-203 expression and its promoter activity (Figures 4B & 4C). These results strongly suggest that BIR domains promote miR-203 transcription. It is reported that promoter region demethylation is involved in upregulation of miR-203 [33]. Thus, we tested whether promotion of miR-203 transcription was due to the regulatory effect of BIR domain on demethylation of miR-203 differentially methylated region (DMR). As shown in Figure 4D, there was no any observable alteration of methylation and unmethylation between T24T cells with either XIAP knockdown cells, or ΔRING domain overexpressed cells, in comparison to parental T24T cells, suggesting that promotion of miR-203 transcription by BIR domains is not through affecting demethylation of miR-203 promoter region. We next bioinformatically analyzed the potential transcription factor binding sites in miR-203 promoter region, and the results revealed that the promoter contains binding sites for multiple transcription factors, including c-Jun, E2F1, Sp1, ELK1, and NF-κB (Figure 4E). The effect of XIAP on these transcription factor expressions was explored in T24T (nonsense) and T24T (shXIAP) cells. The results indicated that knockdown of XIAP in T24T cells only attenuated E2F1 expression, no effect on Sp1 expression and increased p65 and c-Jun (Figure 4F). However, depletion of XIAP expression in T24T cells dramatically inhibited both E2F1‐ and Sp1-dependent transcription activity, which could be completely reversed by ectopic expression of ΔRING domain (Figures 4G & 4H). These results suggest that Inhibition of E2F1‐ and Sp1-dependent transcription activity might be associated with BIR domain attenuation of miR-203 transcription.

**Figure 4.**
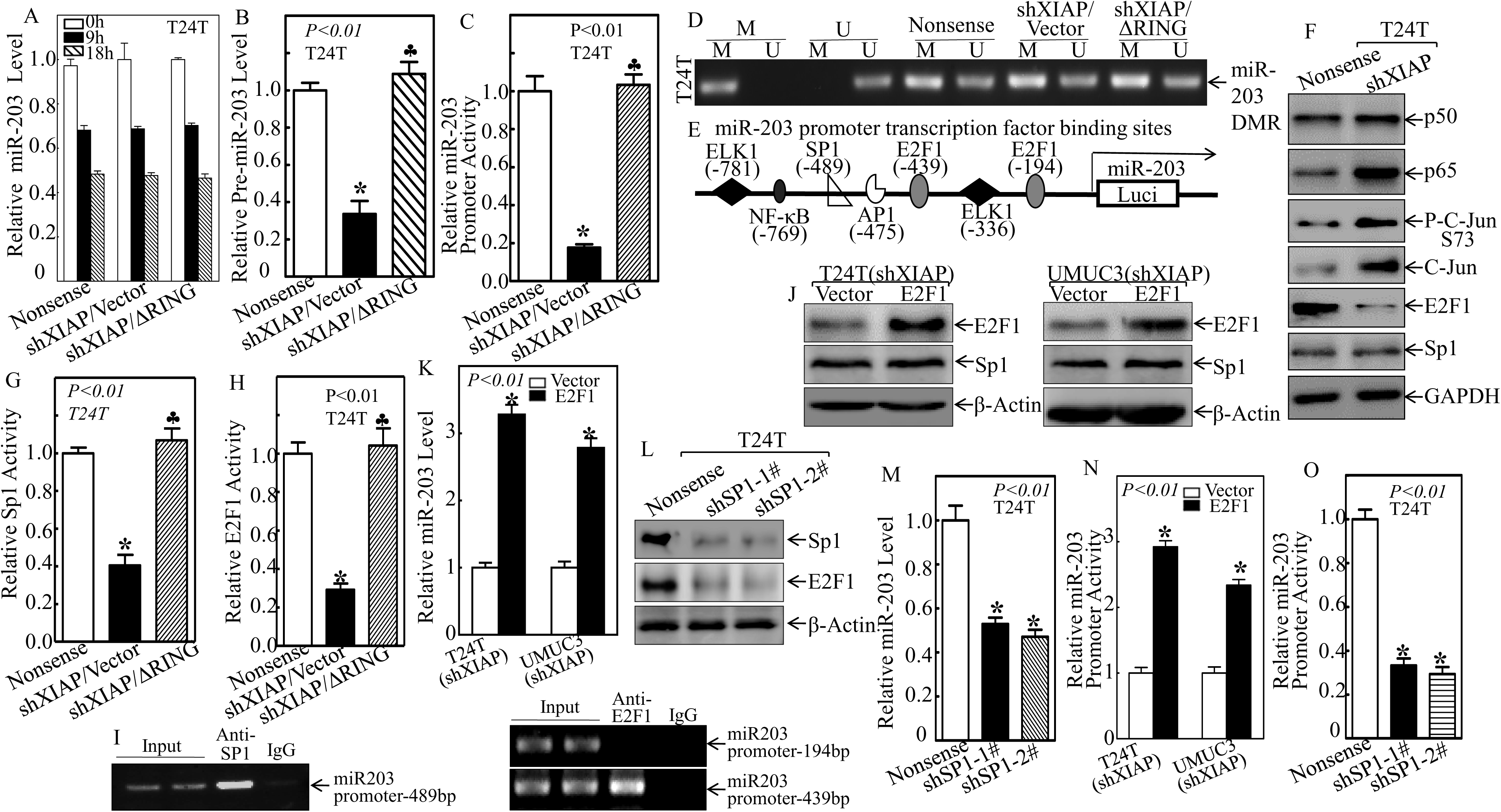
XIAP BIR domains promote miR-203 transcription through transactivation of E2F1 and Sp1 in human BC cells. **(A)** The indicated cells were incubated with actinomycin D (20 μg/ml) for the indicated time periods. Total RNA was isolated, and quantitative real-time PCR was then performed to determine miR-203 levels. The fold change was normalized using GAPDH as the internal control. **(B)** The relative expression levels of pre-miR203 were evaluated with quantitative real-time PCR in the indicated cells. **(C)** The indicated cells were stably transfected with a miR-203 promoter-driven luciferase reporter to determine the miR-203 promoter transcriptional activity. **(D)** XIAP and its BIR domains did not affect miR-203 promoter methylation. **(E)** Schematic representation of the transcription factor binding sites in the human miR-203 promoter-driven luciferase reporter. **(F)** The indicated cell extracts were subjected to Western blot to determinate the functional transcription factors, and GAPDH was used as a protein loading control. **(G and H)** T24T (Nonsense), T24T (shXIAP/Vector), and T24T (shXIAPMRING) cells were transfected with an Sp1-dependent luciferase reporter (G) or an E2F1-dependent luciferase reporter (H), together with pRL-TK. The results are presented as luciferase activity relative to that of vector control transfectants. **(I)** ChIP assay was performed using anti-E2F1 or anti-Sp1 antibody to detect the interaction between E2F1 or Sp1 and the miR-203 promoter. **(J and K)** T24T and UMUC3 cells stably transfected with E2F1 overexpression construct, and the stable transfectants were identified and determined Sp1 expression (J). (K) The stable transfectants were subjected to evaluate the level of miR-203 with real-time PCR, and the symbol (*) indicates a significant increase in miR-203 expression in E2F1 overexpression cells compared with vector transfectants (p < 0.01). **(L and M)** T24T cells were stably transfected with two Sp1 knockdown plasmids separately, and Western blot was employed to determine Sp1 protein expression, and Real-time PCR was performed to determine the miR-203 expression in the stable Sp1 knockdown cells. **(N)** The stable E2F1 overexpression BC cells were stably transfected with a miR-203 promoter-driven luciferase reporter to determine the miR-203 promoter transcriptional activity. **(O)** The stable Sp1 knockdown cells were stably transfected with a miR-203 promoter-driven luciferase reporter to determine the miR-203 promoter transcriptional activity.

To gain direct evidence for the transactivation of the miR-203 promoter by Sp1 and E2F1, a chromatin immunoprecipitation (ChIP) assay was employed to test the potential directly interaction of Sp1 and E2F1 to their putative binding sites in miR-203 promoter region. As shown in Figure 4I, Sp1 did show its binding activity to miR-203 promoter at −489bp, whereas E2F1 was only be able to bind at site −439bp, but not putative binding site at −194 bp of the miR-203 promoter region (Figure 4I). Moreover, overexpression of E2F1 remarkably increased miR-203 expression but did not affect Sp1 expression in both T24T (shXIAP/vector) and UMUC3 (shXIAP/vector) cells (Figures 4J & 4K), while knockdown of Sp1 not only attenuated miR-203 expression activity in T24T cells (Figure 4L & 4M), but also inhibited E2F1 protein expression in T24T cells (Figure 4L). Consistent with E2F1 and Sp1 promotion of miR-203 transcription, ectopic expression of E2F1 in XIAP knockdown cells increased miR-203 promoter activity (Figure 4N), while knockdown of Sp1 significantly decreased miR-203 promoter activity (Figure 4O). These results reveal that both Sp1 and E2F1 play role in XIAP promotion of miR-203 transcription and induction in human BC cells, and Sp1 also acts as upstream regulator of E2F1 to form a positive loop for promoting miR-203 transcription in addition to its direct regulation of miR-203 induction.

### Sp1 is crucial for BIR domains promoting E2F1 transcription and BC cell invasion

We further examined the mechanism of E2F1 upregulation by XIAP. The results showed that knockdown of XIAP expression greatly reduced the mRNA level of E2F1 in both T24T and UMUC3 cells (Figures 5A and 5B). Moreover, depletion of XIAP or Sp1 greatly decreased E2F1 promoter activity, whereas the reduction on E2F1 promoter activity could completely reversed by ectopic expression of ΔRING domain (Figure 5C), strongly indicating that XIAP BIR domains upregulate E2F1 at the transcription level in Sp1 dependent manner. The Sp1 promotion of E2F1 transcription was also supported by the result from a bioinformatics analysis showing that there are three potential binding sites for Sp1 in the E2F1 promoter region (Figure 5D). Consistent with crucial roles of Sp1 and E2F1 in modulation of miR-203 transcription, overexpression of E2F1 in T24T (shXIAP) cells markedly increased the cancer cell invasion, while knockdown of Sp1 in T24T cells significantly inhibited the cancer cell invasion (Figure 5E-5H).

**Figure 5.**
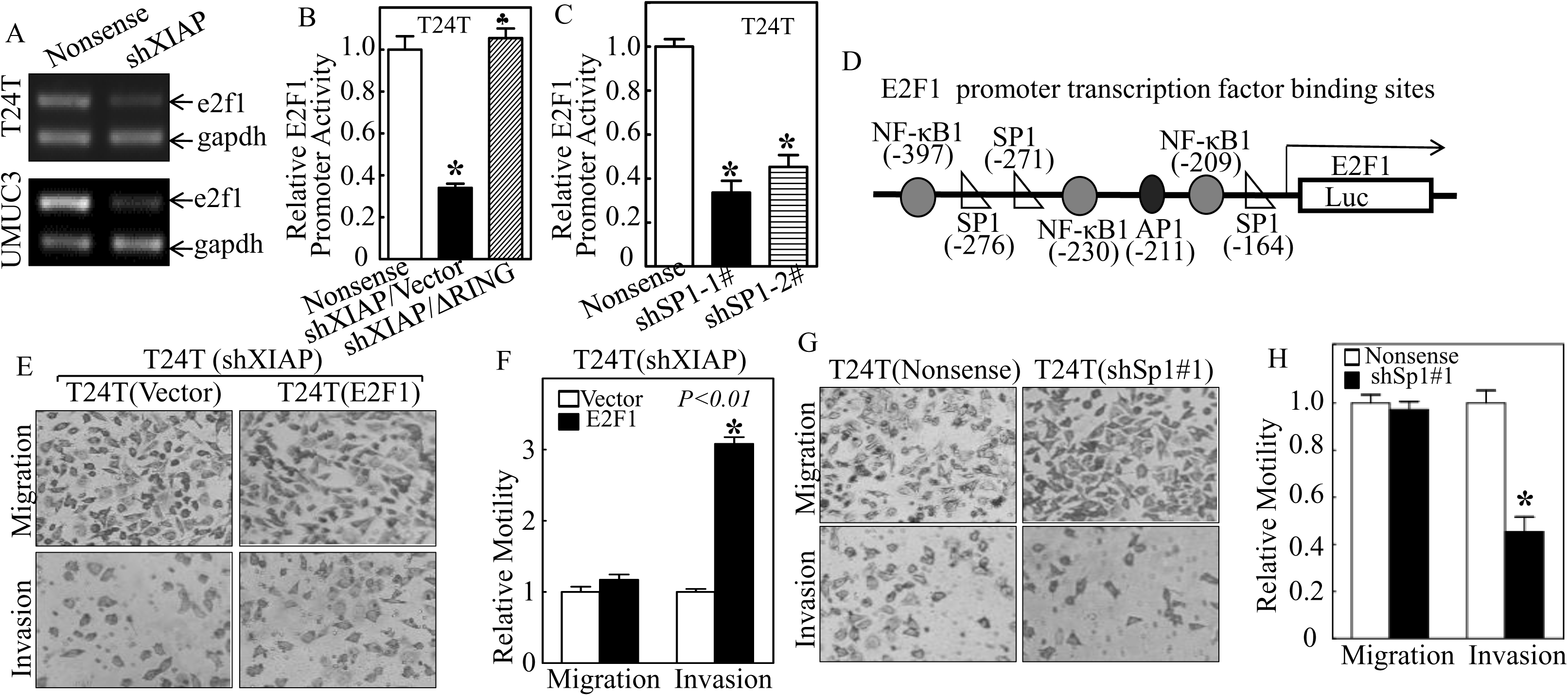
Sp1 is crucial for XIAP BIR domains-mediated E2F1 transcription and promoting BC cell invasion. **(A)** RT-PCR was performed to determine the e2f1 mRNA levels in the indicated cells. **(B)** T24T (Nonsense), T24T (shXIAP/Vector), and T24T (shXIAPMRING) cells were co-transfected with an E2F1 promoter-driven luciferase reporter together with pRL-TK. Then, 24 hours post transfection, the transfectants were extracted to evaluate luciferase activity, with normalization to TK. The results were presented as luciferase activity relative to scramble nonsense transfectant. Each bar indicates the mean±SD of three independent experiments. The symbol (*) and (♣) indicates a significant difference as compared with the vehicle control and T24T (shXIAP/vector), separately (p < 0.05). **(C)** The stable Sp1 knockdown cells were stably transfected with E2F1 promoter-driven luciferase reporter to determine the E2F1 promoter transcriptional activity. **(D)** The potential transcription factor binding sites in the e2f1 promoter. **(E-H)** Different type of transfectants were subjected to cell invasion and migration assays with transwell invasion assay system (E and G). The migration and invasion rates were normalized with the insert control according to the manufacturer’s instructions, and the results are presented as the number of migratory or invasive cells relative to vector control transfectants (F and H).

### BIR2 and BIR3 specifically interacted with E2F1 and Sp1, respectively, to coordinately promote BC invasion

To further elucidate the mechanism of XIAP regulation of Sp1 and E2F1 transcriptional activity, we tested the possibility that the XIAP interacts with Sp1. The results from immunoprecipitation (IP) assay by using T24T cells that expressed HA-XIAP with or without GFP-Sp1. Intriguingly, HA-tagged XIAP and E2F1 was present in the immunoprecipitates following anti-GFP antibodies pulling down of GFP-tagged Sp1 (Figure 6A). This physical interaction was further demonstrated in the immunoprecipitates using anti-HA antibodies pulling down of HA-XIAP (Figure 6B). More interesting is that both Sp1 and E2F1 proteins were present in the co-precipitated protein complex in T24T cells in the absence of XIAP RING domain but not detectable in T24T cells in the absence of XIAP BIR domains (Figure 6C), indicating that XIAP interacts with Sp1 and E2F1 through BIR domains in BC cells. Since that XIAP contains three BIR domains, we further determine whether Sp1 or E2F1 interacts with XIAP through specific BIR domain. The results from co-immunoprecipitation assays using anti-HA antibodies demonstrated that BIR2 domain specially interacted with E2F1, while Sp1 specifically bound to BIR3 domain (Figure 6D). To further investigate the physiological consequence of this physical interaction between Sp1 and XIAP or E2F1 and XIAP in cells upon serum stimulation, we incubated T24T(HA-XIAP) cells in medium containing 20% FBS for 30 min, and the cell extracts were used to perform co-immunoprecipitation assay to pull down endogenous Sp1 and E2F1 by using anti-HA antibodies. The results showed that serum stimulation led to a substantial decrease in XIAP interaction with both Sp1 or E2F1 proteins in BC cells (Figure 6E). Given that pre-miR-203 transcription occurs in the nucleus, we anticipated that the serum stimulation might result in dissociation of XIAP from E2F1 and Sp1 in BC cells. To test this notion, cytoplasmic and nuclear fractions from T24T cells upon serum stimulation were isolated and further subjected to immunoblotting analysis. As shown in Figure 6F, nuclear XIAP translocated to the cytoplasmic upon 20% FBS stimulation, but E2F1 and Sp1 still stayed in nuclear, indicating that nuclear XIAP are mainly responsible for XIAP interaction with Sp1 and E2F1. Furthermore, ectopic expression of BIR2 and BIR3 showed restoration of Src inhibition and MMP2 activation (Figure 6G) and rescued invasion ability (Figures 6H and 6I) in XIAP-deletion BC cells. Consistent with BIR3 promotion of E2F1 transcription *via* Sp1, only ectopic expression of BIR3, but not BIR2, rescued E2F1 protein expression (Figure 6G). Given that our published study indicates the inhibition of Rac1 expression by XIAP [34], it is interesting to define which BIR domain is associated with this function. The results revealed that Rac1 upregulation in XIAP-deficient cells could be specifically abolished by ectopic expression of BIR1, but not either BIR2 or BIR3 (Figure 6G), suggesting that BIR1 mediates XIAP inhibition of Rac1 expression. Consistent with activation of MMP2 by BIR2 and BIR3, ectopic expression of BIR2 and BIR3 also restored E2F1‐ and Sp1-dependent transactivation (Figures 6J and 6K), miR-203 promoter activation, as well as miR-203 expression (Figures 6L and 6M). These results demonstrate that the crosstalk of BIR2 and BIR3 by interaction with E2F1 and Sp1 are drive force for activation of E2F1 and Sp1 leading to miR-203 transcription, Src protein translation inhibition, MMP2 activation and BC invasion.

**Figure 6.**
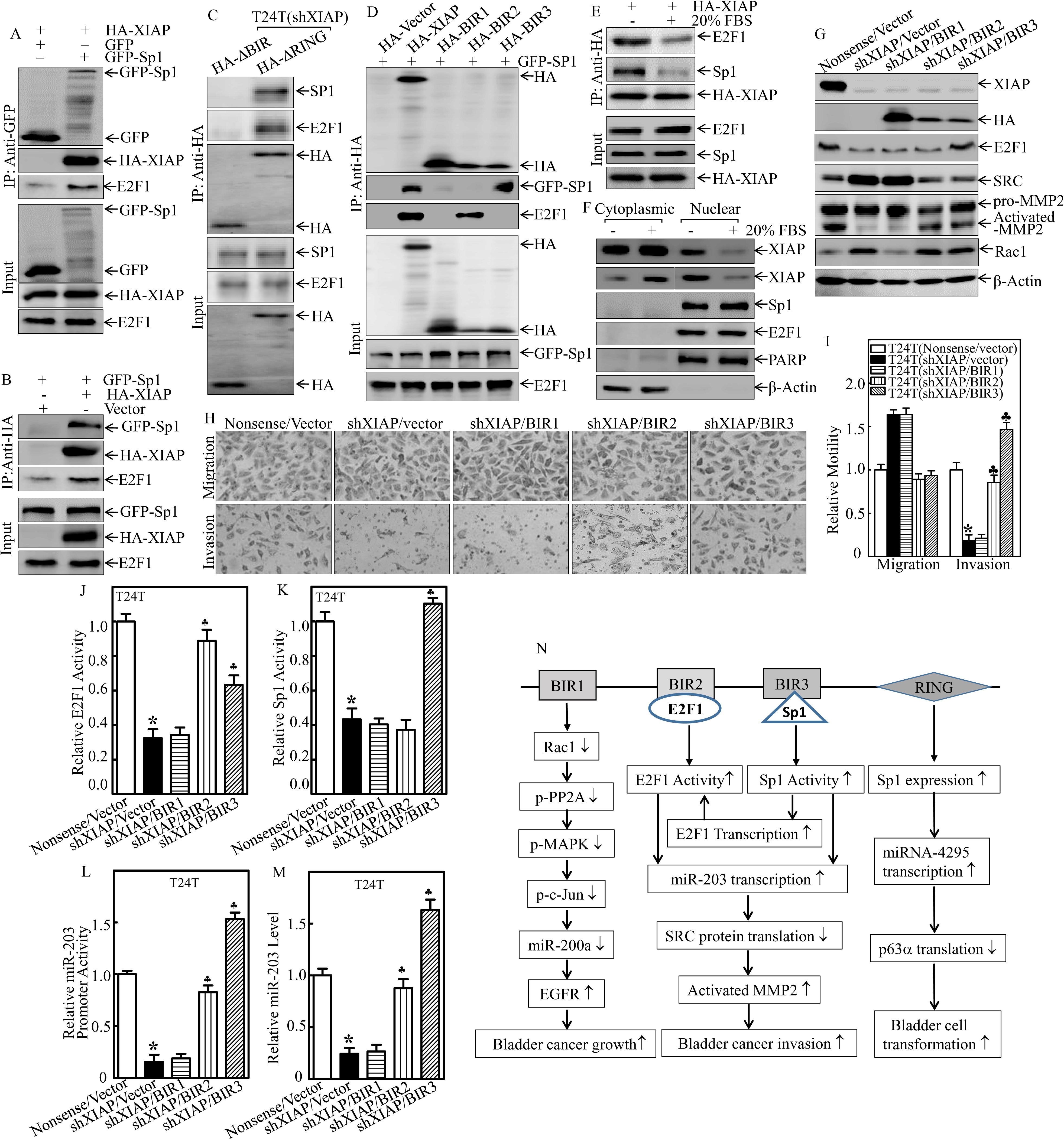
BIR2 and BIR3 domains of XIAP differentially interacted with Sp1 and E2F1 and promote BC invasion. **(A)** Immunoblotting analysis of whole-cell lysates (Input) and GFP-immunoprecipitates (IP) obtained from T24T cells transfected with HA-XIAP with or without combination with GFP-Sp1 using anti-GFP and anti-E2F1 antibody-conjugated beads. **(B)** Immunoblotting analysis of whole-cell lysates (Input) and HA-immunoprecipitates (IP) obtained from T24T cells transfected with GFP-Sp1 alone or in combination with HA-XIAP using anti-HA and anti-E2F1 antibody-conjugated beads. **(C)** Total cellular protein was extracted from the indicated cells, and a co-immunoprecipitation assay was performed using anti-HA antibody-conjugated beads. Immunoprecipitated protein was then subjected to Western blotting to detect the interaction of XIAP BIR domains with antibodies as indicated. **(D)** Immunoblotting analysis of whole-cell lysates (Input) and HA-immunoprecipitates (IP) obtained from T24T cells transfected with GFP-Sp1 with or without combination with various XIAP fragments using anti-HA antibody-conjugated beads. **(E)** Immunoblotting analysis of whole-cell lysates and anti-HA-immunoprecipitates (IP) obtained from T24T(HA-XAP) cells following synchronization overnight in 0.1% fetal bovine serum (FBS) medium and further stimulation with 20% FBS medium for 30 min. **(F)** Immunoblotting analysis of cytoplasmic and nuclear fractions of T24T cells following 24 h of serum deprivation and further stimulation with 20% FBS for 30 min. β-Actin and poly-(ADP-ribose) polymerase (PARP) are cytoplasmic and nuclear markers, respectively. **(G)** The indicated cell extracts were subjected to Western blotting for determination of the expression of XIAP, E2F1, Src and pro-MMP2 and cleaved-MMP2 (activated-MMP2). β-Actin was used as the protein loading control. **(H-I)** Different types of transfectant were subjected to cell invasion and migration assay by using transwell invasion assay system (H). The migration and invasion rates were normalized with the insert control according to the manufacturer’s instructions, and the symbol (*) and (♣) indicates a significant difference compared with the vehicle control and T24T (shXIAP/vector), separately (p < 0.05) (I). **(J and K)** T24T (Nonsense), T24T (shXIAP/Vector), T24T (shXIAP/BIR1), T24T (shXIAP/BIR2) and T24T (shXIAP/BIR3) cells were transfected with an E2F1-dependent luciferase reporter (J) or a Sp1-dependent luciferase reporter (K), together with pRL-TK. The results are presented as luciferase activity relative to that of vector control transfectants. **(L)** The indicated cells were stably transfected with a miR-203 promoter-driven luciferase reporter to determine the miR-203 promoter transcriptional activity. The symbol (*) indicates a significant decrease as compared with nonsense cells as indicated (p < 0.01), while the symbol (♣) indicates a significant increase as compared with T24T (shXIAP/Vector) cells. **(M)** The level of miR-203 in the indicated cells was evaluated with quantitative real-time PCR. **(N)** The schematic of the potential XIAP BIR domains‐ and RING domain-mediated regulation of bladder cancer promotion, and invasion.

## Discussion

Our current study discovered that the BIR domains of XIAP is one of the major factors that promote human BC cell invasion. This important function of XIAP/BIR domains is mediated *via* specific activating MMP2 in Src protein inhibition-dependent manner. Further studies revealed that the BIR domains initiate Sp1‐ and E2F1-mediated transcription of miR-203, which is able to bind the 3’-UTR of src mRNA and ultimately to block Src protein translation. We also identified that activated Sp1 by BIR3 also acts as transcription factor to positively regulation of E2F1 transcription. Most interestingly, the BIR2 is found to specific bind to E2F1, while BIR3 interacts with Sp1 in intact BC cells, and those protein-protein interactions as well as SP1 positively modulation of E2F1 result in ultimately activation of E2F1 and Sp1, and in turn lead to miR-203 transcription, Src protein translation inhibition, and consequently activate MMP2 and BC invasion. This novel mechanistic discovery of BIR2 and BIR3 domains of XIAP in human BC invasion provides highly significant insight into understanding of XIAP in BC invasion.

Matrix metalloproteinases-2 (MMP2) belongs to one of the gelatinases that are primary subgroups of MMPs on the premise of domain structure [35]. MMP2 has a well-known role in degradation of connective tissue stroma and basement membranes, and is a good candidate for a biological marker in many cancers [36]. MMP2 is secreted into the matrix as pro-MMP2 with an auto-inhibitory N-terminal pro-domain [37]. The cysteine switch motif in this domain blocks the catalytic zinc, preventing hydrolysis of substrates [38]. Pro-MMP2 could be activated *via* proteolytic cleavage or chemical disruption of the pro-domain to expose the catalytic zinc for fully enzyme activity [39]. MMP2 activation has been reported to be directly correlated with the aggressiveness of bladder tumors [40]. Here, we are the first to unravel a novel function for XIAP in modulating MMP2 activation that is negatively regulated by proto-oncogene tyrosine-protein kinase Src. We also demonstrate that the crosstalk between BIR2 and BIR3 domains is crucial to inhibit Src protein translation through directly targeting of its mRNA 3’UTR by upregulated miR-203, while the direct protein-protein interactions of BIR2 and BIR3 with E2F1 and Sp1, together with crosstalk between Sp1 and E2F1 result in their strong binding to promoter region of miR-203 leading to its fully transcription, Src inhibition and MMP2 activation in human BC cells. These findings establish a new bridge between XIAP overexpression and MMP2 activation in human BC high invasion, and further help us better understanding XIAP induction associated with the progression and aggression of malignant bladder tumor development [14,15]. Further investigation will be mainly focusing on the precise role of XIAP *in vivo* by using XIAP knockout, or overexpression, as well as each BIR domain deletion knock-in mouse model.

The function of Src in cancer biology in general is dependent on the cancer types. For example, Src is overexpressed or activated in breast, prostate, colorectal, pancreatic, hepatocellular, esophageal, head and neck, ovarian, and lung cancer as well as in leukemia and lymphoma [41]. Src is the oldest and best-studied proto-oncogene, and its high expression or activation is positively associated with tumor grade and stage in these cancers [42-44]. However, the divergence of the expression and activity of Src are reported in bladder cancers. It has been reported that the Src expression level and activity are surprisingly low in human BC cell lines including TccSup, T24, and U5637 [45]. Moreover, compared with high-grade counterparts, low-grade BC cell lines and tumors possess high expression and activity of Src [46]. Src protein levels are also attenuated with increasing bladder cancer stage and affected cancer metastasis [28,47]. Our current study revealed low mRNA and protein expression of Src in human and mouse BC tissues. Thus, the studies from our laboratory and others suggest that Src is a potential tumor suppressor in BCs. To the best of our knowledge, we unprecedented discover that Src-mediated BC invasion mainly rely on its downstream powerful BC invasion/metastatic effector, MMP2. We also demonstrate that Src-associated MMP2 regulation only target MMP2 activation rather than its expression in BIR domains of XIAP-dependent. These new findings not only help us be more aware of the tumor suppressive role of Src in BCs, but also warn us to consider the tissue-specificity of drugs targeting Src. Further study need to be elucidated about how Src-regulated MMP2 activation in our future endeavors.

It has previously been reported that miR-203 is significantly upregulated in bladder cancers [48], indicating that it may function as a tumor promoter in the disease progression. In the present studies, we found that miR-203 had an essential role in XIAP regulation of Src expression *via* binding to 3’-UTR of Src mRNA. Consistent with its oncogenic role of XIAP in Src protein expression, we found that miR-203 significantly inhibited Src protein translation without affecting its mRNA. Furthermore, we showed that XIAP promoted miR-miR-203 expression through enhancing Sp1 and E2F1 activation. Sp1 and E2F1 are both transcription factors, and their expression and activity has been reported to be elevated in many cancers [49,50]. We reported here that XIAP might act as a promising natural promoter of Sp1 and E2F1 through specially interacting with them and enhancing their activity. Moreover, we found that Sp1 or E2F1 was mainly bound to nuclear XIAP in unstimulated BC cells, but dissociated with XIAP resulting from nuclear XIAP shuttled to cytoplasm following serum stimulation. Additionally, the molecular mechanism that mediates the dissociation of Sp1 and E2F1 from XIAP in BC cells upon serum stimulation also merits further investigation. Our previous report reveals that in HCT116 colon cancer cells, the BIR domains of XIAP could bind E2F1 to promote cell growth by strengthening cyclin E expression [21]. Our current finding further extends this knowledge revealing that E2F1 binds to XIAP BIR2 domain and made a new discovery of Sp1 interaction with XIAP BIR3 domain, and Sp1 also acts as an E2F1 upstream transcriptional factor to initiate a crosstalk with E2F1 to promote BC cell invasion. The crosstalk between BIR2 and BIR3 domains in regulation of miR-203/SRC/MMP2 axis is greatly supported by the findings that ectopic expression of either BIR2 or BIR3 could restore E2F1-dependent transactivity, whereas only BIR3, but not BIR2, rescued Sp1-dependent transactivity. Consistently, the defect of miR-203 promoter-driven reporter activity and miR-203 expression in XIAP knockdown cells could be completely restored by ectopic expression of either BIR2 or BIR3. These results suggesting that Sp1 acts as an E2F1 upstream regulator for XIAP promotion of MMP2 activation and invasion of BC cells.

In summary, our studies have revealed a novel Sp1/E2F1/miR-203/Src pathway that is responsible for activation of MMP2 and the tumor-promotive role of XIAP in BC cell invasion. We show a new link between XIAP, Src and MMP2 activation, which may be BC specificity. More importantly, we identify two physical protein-protein interactions: XIAP and Sp1, XIAP and E2F1, and further point out that BIR2 domain of XIAP is essential and sufficient for its interaction with E2F1, while BIR3 domain of XIAP is mainly responsible for its binding with Sp1. In addition, we find that BIR3 of XIAP-initiated Sp1 also acts as an upstream regulator for E2F1 transcription. Collectively, our findings from current studies, for the first time to the best of our knowledge, demonstrate an essential role of crosstalk between BIR2 and BIR3 of XIAP by their interactions with E2F1 and Sp1, respectively, to activate MMP2 and BC invasion by inhibiting Src protein translation in miR-203-dependent manner. Our findings provide novel molecular evidence that contributes to an improved understanding of the tumor-suppressive role of Src and its relationship with the BIR domain of XIAP and MMP2 activation in bladder cancer cells, suggesting that Src or the BIR domains of XIAP could potentially be used as a therapeutic target in future BC therapy. Finally, these findings also provide a clue for us to understand the reason that high nuclear expression of XIAP is associated with poor clinical outcome of cancer patients that is observed in clinical studies [51].

## Materials and methods

### Cell lines, plasmids, antibodies, and other reagents

The human invasive BC cell line UMUC3 was provided by Dr. Xue-Ru Wu (Department of Urology and Pathology, New York University School of Medicine, New York, NY), and was used in our previous studies [52]. The human metastatic BC cell line T24T, which is a lineage-related metastatic lung variant of the invasive BC cell line T24 [53], was kindly provided by Dr. Dan Theodorescu [54] and used in our previous studies [55]. UMUC3 cells were maintained in Dulbecco’s modified Eagle’s medium (DMEM) supplemented with 10% FBS (HyClone, Logan, UT), 1% penicillin/streptomycin and 2 mM L-glutamine (Life Technologies, Rockville, MD). T24T cells were cultured in DMEM/Ham's F-12 (1:1 volume) mixed medium supplemented with 5% FBS, 1% penicillin/streptomycin and 2 mM L-glutamine. The shRNA that specifically targets human XIAP and Sp1 was purchased from Open Biosystems (GE, Pittsburgh, PA). HA-ΔBIR and HA-ΔRING expression plasmids were described in our previously studies [20,21,25]. miR-203 mimic RNA was kindly provided by Dr. Dale D. Tang (The Center for Cardiovascular Sciences, Albany Medical College, Albany, New York) [56]. The Src expression plasmid was obtained from Addgene (Cambridge, MA). E2F1‐ and Sp1-dependent luciferase reporters were described in our previous papers [25,57]. The human Src mRNA 3’-UTR luciferase reporters and its mutant (the binding site of miR-203 was mutated) were cloned into a pMIR-report luciferase vector. The plasmid containing the luciferase reporter under control of human miR-203 gene promoter was constructed into a PGL3-BASIC vector. Anti-XIAP antibody was purchased from Becton, Dickinson and Company (Franklin Lakes, NJ). Specific antibodies against HA, Src, S6 ribosomal protein, P-S6 ribosomal protein Ser235/236, p53, c-Jun, P-c-Jun at Ser73, NF-κB p65 and GAPDH were purchased from Cell Signaling Technology (Beverly, MA). Antibodies specific for Sp1, E2F1, MMP2 and β-Actin, were bought from Santa Cruz (Dallas, TX). Antibodies specific against p50 were bought from Abcam (Cambridge, MA, USA). The protein synthesis inhibitor cycloheximide (CHX) was purchased from Calbiochem (San Diego, CA, USA). The dual luciferase assay kit was purchased from Promega (Madison, WI, USA). TRIzol reagent and the SuperScript™ First-Strand Synthesis system were bought from Invitrogen (Grand Island, NY, USA). PolyJet™ DNA In Vitro Transfection Reagent was purchased from SignaGen Laboratories (Rockville, MD, USA). Both the miRNeasy Mini Kit and the miScript PCR system for miRNA detection were bought from Qiagen (Valencia, CA, USA).

### Human bladder cancer tissue samples

Twenty pairs of primary bladder cancer samples and their paired adjacent normal bladder tissues were obtained from patients who underwent radical cystectomy at the Department of Urology of the Union Hospital of Tongji Medical College (Wuhan, China) between 2012 and 2013. All specimens were immediately snap-frozen in liquid nitrogen after surgical removal. Histological and pathological diagnoses were confirmed, and the specimens were classified by a certified clinical pathologist according to the 2004 World Health Organization Consensus Classification and Staging System for bladder neoplasms. All specimens were obtained with appropriate informed consent from the patients, and a supportive grant was obtained from the Medical Ethics Committee of China. The experiments were carried out in accordance with The Code of Ethics of the World Medical Association (Declaration of Helsinki) for experiments involving human studies.

### Animal experiments and immunohistochemistry-paraffin (IHC-P)

Male C57BL/6J mice at the age of 5~6 weeks were randomly divided into two groups, with 12 mice in each group, including a vehicle-treated control group and an N-butyl-N-(4-hydroxybutyl) nitrosamine (BBN)-treated group. Mice in the BBN-treated group received BBN (0.05%) in drinking water for 20 weeks, while vehicle-treated group was provided with normal drinking water containing same amount of DMSO. The mice were sacrificed at the end of the experiment and mouse bladder tissues were excised and fixed overnight in 4% paraformaldehyde at 4°C. Fixed tissues were processed for paraffin embedding, and the serial 5-μm-thick sections were then immunostained with specific antibodies against Src (Cell Signaling Technology). The resultant immunostaining images were captured using an AxioVision Rel.4.6 computerized image analysis system (Carl Zeiss, Oberkochen, Germany). Protein expression levels were presented by the integrated optical density per stained area (IOD/area) that was analyzed with Image-Pro Plus version 6.0 (Media Cybernetics, MD). Briefly, the IHC stained sections were evaluated at 400-fold magnifications, and at least 5 representative staining fields in each section were analyzed to calculate the optical density based on typical images that had been captured.

### Western Blot

Western Blot were assessed as previously described [58]. Briefly, cells were plated in 6-well plates and cultured in normal FBS medium until 70–80% confluent. The cells were then cultured in 0.1% FBS medium for 12 hours, followed by treatment with different doses of ISO for the indicated time. The cells were washed once with ice-cold phosphate-buffered saline, and cell lysates were prepared with a lysis buffer (10 mM Tris-HCl (pH 7.4), 1% SDS, and 1 mM Na3VO4). An equal amount (80 μg) of total protein from each cell lysate was subjected to Western blotting with the indicated antibody. Immunoreactive bands were detected using alkaline phosphatase-linked secondary antibody and an ECF Western blotting system (Amersham Biosciences, Piscataway, NJ). Images were acquired using a Typhoon FLA 7000 imager (GE Healthcare, Pittsburgh, PA).

### RT-PCR and quantitative RT-PCR

Total RNA was extracted with TRIzol reagent (Invitrogen Corp. USA), and cDNAs were synthesized with a SuperScript III First-Strand Synthesis System for RT-PCR (Invitrogen Corp. USA). A pair of oligonucleotides (Forward: 5’-GATGATCTTGAGGCTGTTGTC-3’ and Reverse: 5’-CAGGGCTGCTTTTAACTCTG-3’) were used to amplify human GAPDH cDNA as a loading control. The human Src cDNA fragments were amplified with a pair of human Src-specific PCR primers (Forward: 5’-TCCGACTCCATCCAGGCTGA-3’ and Reverse: 5’-TGTCCAGCTTGCGGATCTTG-3’). The human E2F1 cDNA fragments were amplified with 5’-GAGGTGCTGAAGGTGCAGAA-3’; (Forward) and 5’-GTTTGCTCTTAAGGGAGATCTG-3’ (Reverse). The PCR products were separated on 2% agarose gels, stained with ethidium bromide (Fisher Scientific Corporation, USA), and scanned for imaging under UV light. The results were visualized with a Alpha Innotech SP Imaging System (Alpha Innotech Corporation, San Leandro, CA, USA). Quantitative RT-PCR was performed to examine the expression level of mature miRNAs and pre-miRNA, as described previously [59].

### [^35^S] Methionine pulse new protein synthesis assays

Cells were incubated with methionine-cysteine free DMEM (Gibco-BRL, Grand Island, NY, USA) containing 2% dialyzed fetal calf serum (Gibco-BRL) and 10 μM MG132 for 30 minutes and then incubated with 2% FBS methionine-cysteine-free DMEM containing 35S-labeled methionine/cysteine (250 μCi per dish, Biomedicals, Inc., Irvine, CA) for the indicated periods. The cells were extracted with lysis buffer (Cell Signaling Technology) containing a complete protein inhibitor mixture (Roche) on ice, and 500 mg of total lysate was incubated with anti-Cyclin D1 antibody-conjugated agarose beads (R&D Systems, Minneapolis, MN, USA) overnight at 4°C. The immunoprecipitates were washed with the cell lysis buffer five times, heated at 100°C for 5 min and subjected to sodium dodecyl sulfate polyacrylamide gel electrophoresis. The membranes were then subjected to autoradiography for determination of the newly synthesized 35S-labeled Cyclin D1 protein as described in our previous studies[60].

### Luciferase assay

T24T and UMUC3 cells were transfected with the indicated luciferase reporter construct in combination with a pRL-TK vector (Promega, Madison, WI). The transfectants were seeded into 96-well plates and cultured for 12 hours. The cells were then extracted with luciferase assay lysis buffer (Promega, Madison, WI) and subjected to determination of luciferase activity using a luciferase assay system (Promega Corp., Madison, WI) with a microplate luminometer LB 96V (Berthold GmbH & Co. KG, Bad Wildbad, Germany). The luciferase activity was normalized to the internal control TK activity based on the manufacturer’s instructions.

### Methylation-specific PCR

Genomic DNA was isolated with a DNeasy Blood & Tissue Kit (Qiagen) according to the manufacturer's instructions. Genomic DNA (2 mg) was treated with sodium bisulfite using an EpiTect Bisulfite Kit (Qiagen). Methylation-specific PCR was performed using 20 ng of bisulfite-converted DNA and specific primers. Methylated primer and unmethylated primers for the miR-203 promoter at the differentially methylated region (DMR) were designed according to a previous study [61]. PCR products were run on a 2% agarose gel and visualized after ethidium bromide staining. Bisulfite-converted methylated and unmethylated DNA from the EpiTect PCR Control DNA Set (Qiagen) were used as positive and negative controls.

### Immunoprecipitation

For immunoprecipitation experiments, cells transfected with the indicated plasmids were collected and lysed in 1 × Cell Lysis Buffer (Cell Signaling Technology, Danvers, MA, USA) containing protease inhibitors (Roche, Branchburg, NJ, USA) followed by brief sonication. Any insoluble material was removed by centrifugation at 16,000×g for 20 minutes at 4°C. Immunoprecipitation was carried out by incubation of cell lysates with anti-HA or anti-GFP antibody-conjugated agarose beads. After an overnight incubation, beads were washed three times with immunoprecipitation lysis buffer, and bound proteins were subjected to Western Blot assay [59].

### *In vitro* cell migration and invasion assays

*In vitro* migration and invasion assays were conducted using transwell chambers (for migration assays) or transwell chambers pre-coated with Matrigel (for invasion assays) according to the manufacturer's protocol (BD Biosciences, Bedford, MA), as described previously [52]. Briefly, 700 μl of medium containing 10% FBS (for UMUC3 and T24T cells with different transfectants) was added to the lower chambers, while homogeneous single cell suspensions (5×10^4^ cells/well) in 0.1% FBS medium as indicated were added to the upper chambers. The transwell plates were incubated in a 5% CO_2_ incubator at 37°C for 24 hours and thereafter were washed with PBS, fixed with 4% formaldehyde, and stained with Giemsa stain. The non-migrating or non-invading cells were scrapped off the top of the chamber. The migration and invasion rates were quantified by counting the migratory and invasive cells in at least five random fields under a light microscope (Olympus, Center Valley, PA).

### Statistical methods

Associations between categorical variables were assessed using a chi-square test. Student’s t-test was utilized to compare continuous variables, and the results are summarized as the means ± SD between different groups. Paired t-tests were performed to compare the difference between paired tissues in the real-time PCR analysis. p < 0.05 was considered statistically significant.

## Acknowledgement

We thank Dr. Dale D. Tang from Albany Medical College for providing us with miR-203 mimic RNA. This work was partially supported by the grants of NIH/NCI CA165980, CA1776655 and CA112557, and NIH/NIEHS ES000260.

## Author contributions

Conceptualization, J.H.X. and C.S.H.; Methodology, J.X.L., J.H.X., and H.L.J.; Investigation, J.H.X., Z.X.T., X.H.H., J.L.Z., and M.W.H.; Writing – Original Draft, J.H.X. and C.S.H.; Writing – Review & Editing, C.S.H. and H.S.H.; Funding Acquisition, C.S.H.; Resources, H.S.H and C.S.H.; Supervision, R.Y, C.S.H., and H.S.H.

